# Lipidomics profiling identifies β-oxidation as a key process in noise-induced hearing loss

**DOI:** 10.1101/2025.03.25.645361

**Authors:** Gunseli Wallace, Lingchao Ji, Luis R Cassinotti, Maureen Kachman, Costas A Lyssiotis, Charles F Burant, Gabriel Corfas

**Affiliations:** Kresge Hearing Research Institute and Department of Otolaryngology-Head and Neck Surgery, University of Michigan, Ann Arbor, MI, United States; Cellular and Molecular Biology Program, University of Michigan Medical School, Ann Arbor, MI 48109, USA; Department of Internal Medicine, University of Michigan Medical School, Ann Arbor, Michigan 48109, United States; Department of Molecular & Integrative Physiology, University of Michigan, Ann Arbor, MI 48109

## Abstract

Noise-induced hearing loss (NIHL) is the second leading cause of hearing loss worldwide, and the most common cause in young adults. Despite this burden, the molecular mechanisms by which noise causes damage are poorly understood, and there are no pharmacologic therapies to prevent or reduce noise-induced damage to the inner ear. Here, using targeted and untargeted lipidomics, we show that noise exposure induces changes in fatty acid (FA) and acylcarnitine (CAR) species in the inner ear, a metabolic profile indicative of noise-induced increases in β- oxidation. This conclusion is validated through treatment with Etomoxir, an inhibitor of carnitine palmitoyltransferase 1A, the rate-limiting enzyme of long-chain β-oxidation. Furthermore, we demonstrate that blocking β-oxidation with Etomoxir does not affect hearing in a normal acoustic environment but reduces the extent of hearing loss induced by an intense noise exposure (2 hours, 112 dB SPL, 8-16kHz). Together, our findings provide insights into cochlear energy metabolism and suggest that its modulation could be targeted to reduce NIHL.

## Introduction

Hearing loss is a major global health concern, affecting more than 20% of the world’s population. Noise contributes significantly to this burden; one-quarter of hearing loss globally is suggested to be caused by noise overexposure, and more than 15% of U.S. teenagers show evidence of noise-induced hearing loss (NIHL) on audiological testing^1^. The impact on young adults is particularly concerning as hearing loss is often permanent due to a lack of regenerative capacity in the inner ear sensory epithelium, also known as the organ of Corti, and the unavailability of FDA-approved pharmacologic therapies to prevent, reduce, or treat NIHL.

The two types of cochlear sensory hair cell populations and their neural connections have received particular attention in NIHL due to their importance in sound perception^2^. Inner hair cells (IHCs) convert sound-driven vibrations into neural signals through their synapses with spiral ganglion neurons (SGNs), which then send auditory information to the brain through the ascending auditory pathway. Outer hair cells (OHCs), through their electromotility, regulate cochlear amplification. Noise-induced inner ear damage occurs in a stereotypical pattern that depends on the intensity and duration of the exposure. Animal studies indicate that the synapses between the IHCs and the SGNs are the most susceptible structure in the organ of Corti to noise. For example, in mice, a 2-hour exposure to ∼100 decibel sound pressure level (dB SPL), similar to that at a concert or loud sporting event, is sufficient to cause significant IHC-SGN synapse loss, resulting in what is now called hidden hearing loss. Though undetectable in humans using standard audiometry tests, animal studies indicate that this pathology causes defects in temporal auditory processing and hearing in noise^2,3^. Greater exposures cause hair cell damage or loss, which results in permanent and overt hearing loss, i.e., elevated hearing thresholds on audiological testing^4^.

While the impact of noise on inner ear structure and function has been well characterized, the molecular processes initiated by noise exposure that might contribute to NIHL remain poorly understood. To address this gap in knowledge, we previously developed a targeted metabolomics pipeline to interrogate the effects of noise on inner ear metabolism^5^. Our early work, which assessed central carbon metabolism in isolated otic capsules, identified noise-induced alterations in metabolite levels, including changes in several amino acids, molecules involved in oxidative stress (e.g. oxidized glutathione), components of the TCA cycle (e.g. succinate and citrate), and molecules involved in lipid metabolism (e.g. acetyl-l-carnitine). However, the targeted nature of this initial analysis, which focused on water-soluble metabolites, left the potential effects of noise on lipids, which have been speculated to be involved in hearing loss^6–8^ and noise-induced damage^9–11^, unexplored.

Here, using targeted and untargeted lipidomics profiling of otic capsules after exposure at two levels of noise intensity, we identify significant noise-induced changes in fatty acids (FAs), specifically long and very-long unsaturated FAs, as well as alterations in acylcarnitines and free carnitine, suggesting β-oxidation is induced by noise exposures. Furthermore, using Etomoxir, a carnitine palmitoyl transferase 1A (CPT-1A) inhibitor, we demonstrate that prevention of fatty acid oxidation reduces the degree of noise-induced damage, suggesting potential prophylactic strategies to reduce NIHL.

## Results

### Untargeted lipidomics identifies noise-induced changes in the levels of unsaturated fatty acids (FAs) in the otic capsule

In initial studies, mice were exposed either to noise (8-16 kHz at 100 dB SPL for 2 hours) or placed in the noise exposure chamber for the same length of time with the speaker off for controls. Immediately after the exposure, the entire otic capsules were collected and flash frozen. Later, lipids were extracted and subjected to mass spectroscopy-based lipidomics analysis (MS; Figure 1A). After quality control (see Methods), 735 unique lipids from 26 lipid classes were identified. Principal component analysis (PCA) revealed clear separation between noise- and sham-exposed samples (Figure 1B), demonstrating that noise exposure alters otic capsule lipid levels.

**Figure 1:**
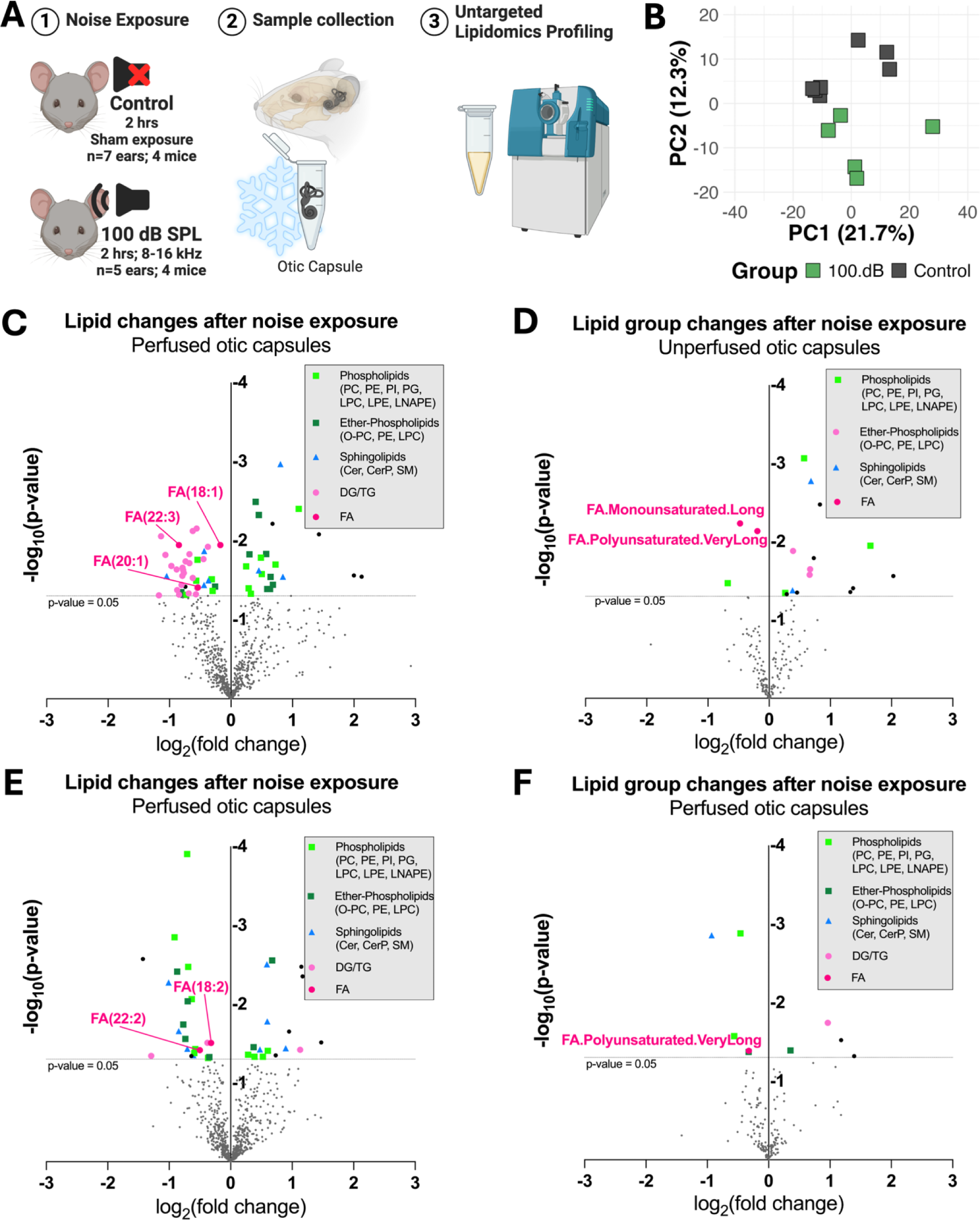
Noise alters otic capsule fatty acid metabolism. **(A)** Experimental design for panels B-D. **(B)** PCA of noise- and sham-exposed otic capsules (untargeted lipidomics) shows clear separation. **(C-D)** Volcano plot showing lipid species (C) or lipid groups (D) whose otic capsule levels are changed by noise exposure in mice without cardiac perfusion. Data represents n=7 (sham), n=5 (noise) otic capsules from 4 mice in each group. **(E-F)** Volcano plot showing lipid species (E) or lipid groups (F) between noise- and sham-exposed mice with cardiac perfusion. Data represents 5 otic capsules from 3 mice per group. In **D** and **F**, the lipid groups were created based on species headgroup, chain length, and saturation. Headgroups: PC, phosphatidylcholine; PE, phosphatidylethanolamine; PI, phosphatidylinositol; PG, phosphatidylglycerol; LPC, lysophosphatidylcholine; LPE, lysophosphatidylethanolamine; LNAPE, N-acyl phosphatidylethanolamine; O-, ether; Cer, ceramide; CerP, ceramide phosphate; SM, sphingomyelin; DG, diacylglycerol; TG, triacylglycerol; FA, fatty acid. Chain Length: Very Long: >20 carbons (C)/side chain (average); Long: 12-20, Medium: 6-12. Saturation: Polyunsaturated: >1 units unsaturation/chain (average), Monounsaturated: 0-1, Saturated: 0. Dots in panels C-F represent either one lipid, or lipid group, and are color-coded when p<0.05 (nominal). Unpaired two-sided Student’s t-test was used for statistical analysis; nominal p = 0.05 is indicated by the dashed horizontal line.

Differential analysis of lipid changes between noise and sham exposed samples identified 22 lipid species that were increased by noise exposure and 39 that were decreased. Lipid species associated with energy, including triglyceride (TG), diglyceride (DG), and fatty acid (FA) species were decreased by noise, whereas several membrane lipids including phospholipids, ether-linked phospholipids, and sphingolipids were increased (Figure 1C). To identify the broader metabolic processes elicited by noise exposure, we grouped the lipids based on saturation, headgroup, and chain length, properties known to influence their energetic, signaling, and membrane function^12–14^. MS intensities within each group were summed and compared between noise and sham-exposed samples. This analysis showed significant decreases in very long chain polyunsaturated FAs and long chain monounsaturated FAs and increases in several phospholipid species (Figure 1D).

The observed concurrent decrease in FA, TG, and DG species suggested that this noise exposure might induce the release of FAs from TGs found in lipid droplets. Since these signals could reflect changes in either systemically circulating or local lipid pools, we repeated the untargeted analysis using mice that underwent cardiac perfusion to remove blood immediately after noise exposure. Like the unperfused samples, very long, polyunsaturated FAs decreased by a similar magnitude and significance in the perfused samples (Figure 1E, F). Together, these results indicate that noise alters FA metabolism in otic capsule tissues, potentially to meet the increased local energetic demands of noise exposure^15,16^.

### Targeted lipidomics reveals noise intensity-dependent changes in otic capsule fatty acid metabolism

Our previous study showed that noise significantly increases the levels of otic capsule acetyl-l- carnitine^5^, a molecule produced during β-oxidation, the catabolic process that breaks down fatty acids for energy. While β-oxidation serves as a critical energy source during cellular stress in many tissues^17^, its role in the inner ear, particularly during noise exposure, remains unexplored. Therefore, to validate the changes in FAs and interrogate potential alterations in β-oxidation, we performed targeted lipidomics profiling of FAs and CARs on otic capsule extracts from mice exposed to the same noise as above (100 dB SPL; 8-16kHz; 2 hours) and sham controls.

In agreement with the untargeted findings, noise exposure caused a decrease in both the very long and long chain polyunsaturated FAs (3 out of 4 and 5 out of 8 significantly changed, respectively), long monounsaturated FAs (4 out of 5), and free carnitine. Reciprocal changes were seen in the CAR species, with significant increases in long polyunsaturated (5 out of 6) and monounsaturated (3 out of 4) species (Figure 2A-C), consistent with the hypothesis that this level of noise increases FA influx into mitochondria where they are utilized for β-oxidation.

**Figure 2:**
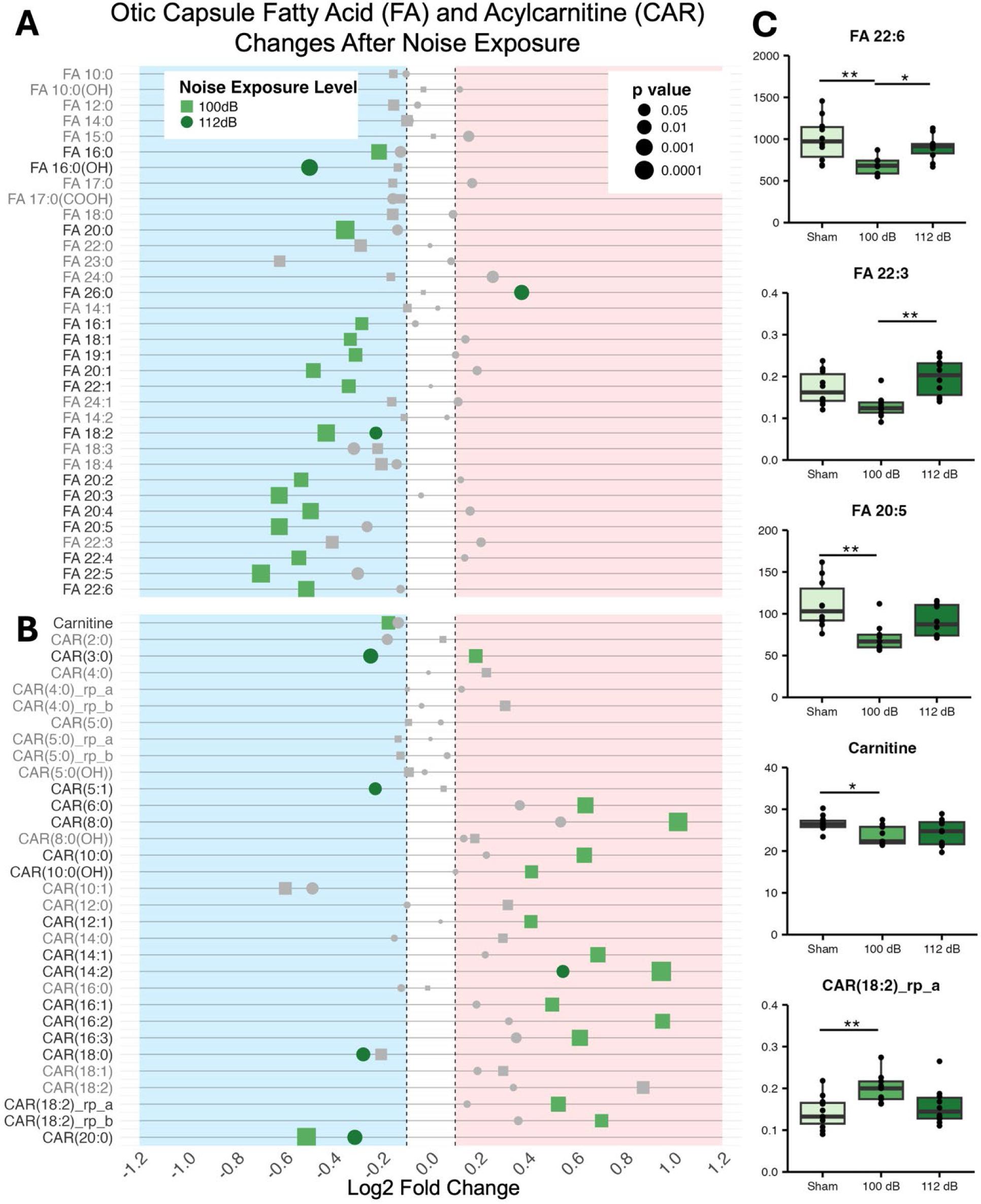
A 2-hour exposure to 100 dB SPL 8-16 kHz noise band, but not to 112 dB SPL, decreases otic capsule fatty acids (FAs) and free carnitine, and increases acylcarnitine (CAR) concentrations, suggestive of increased β-oxidation. **(A-B)** Noise causes significant changes to both fatty acid (FA) and acylcarnitine (CAR) species. Sham-exposed animals were used as controls. Two-way ANOVA with Tukey’s multiple comparison test was used for statistical analysis. Dots represent one lipid species and are color-coded when adjusted p <0.05. The dot’s shape and color correspond to the noise exposure level. **(C)** Quantitative levels of selected lipid species in sham- and noise-exposed samples. Values are in µg (lipid) / mg (sample). Two-way ANOVA with Tukey’s multiple comparison test was used for statistical analysis. Dots represent otic capsule samples. n=10 otic capsules from n=5 mice (sham); n=9 otic capsules from n=5 mice (noise). *p < 0.05; **p < 0.01.

Since noise intensity determines the magnitude of change for many central carbon metabolites^5^, we examined if lipid changes show a similar intensity dependence by repeating the profiling on a cohort of mice exposed to 112 dB SPL (8-16kHz; 2 hours), a level which corresponds to roughly 16 times the acoustic energy imparted by a 100 dB SPL exposure of the same length. Intriguingly, this significantly greater exposure caused only minimal alterations in FAs and CARs (Figure 2A- C). These findings suggest one of two possible scenarios: either β-oxidation is utilized only at lower noise levels, or FA mobilization increases at higher intensities to maintain stable lipid levels in response to continued β-oxidation.

### Noise exposure increases inner-ear FA consumption through β-oxidation

The rate-limiting step of long-chain fatty acid β-oxidation is their transport into the mitochondria by CPT-1A. To directly explore the role of β-oxidation in both normal inner ear function and the response to noise, we inhibited this transfer by daily treatment with Etomoxir, an irreversible inhibitor of CPT-1A^18–20^ (35 mg/kg/day via intraperitoneal (IP) injection).

First, to interrogate β-oxidation activity in the normal acoustic environment, we compared the otic capsule lipidomes of mice treated with Etomoxir or saline daily for 16 days. Otic capsules were harvested several hours after the last injection, and targeted lipid profiling of FAs and CARs was performed. Etomoxir treatment did not affect the very-long chain polyunsaturated FAs that were reduced by a 100 dB SPL noise exposure, but, consistent with inhibition of CPT-1A, caused an accumulation of long unsaturated FAs (Supplemental Figure 1). However, a significant accumulation of polyunsaturated (5 out of 6) and monounsaturated (3 out of 4) CAR species was detected, a lipid change inconsistent with inhibition of β-oxidation. These results suggest β- oxidation activity is not a major contributor to the energy production in the inner ear at rest, and are consistent with previous studies suggesting a preference for aerobic glycolysis and a low metabolic rate in the inner ear under normal acoustic conditions^16,21^.

Next, to directly test the hypothesis that β-oxidation increases during noise, we treated mice with Etomoxir and repeated the targeted FA and acylcarnitine profiling after noise exposure (100 dB SPL or 112 dB SPL) and sham exposure. If the noise-induced FA and CAR changes seen in animals not treated with Etomoxir are indeed caused by increased β-oxidation, we anticipated that in mice with Etomoxir treatment, the effects of noise-exposure on FAs and CARs would be lost or, alternatively, FA levels would increase, indicating mobilization of FA. The results were consistent with these predictions, though there were quantitative differences between the two exposure levels. Etomoxir treatment abolished the effects of the 100 dB SPL exposure on FA and CAR changes. In contrast, the 112 dB SPL exposure resulted in accumulation of very long and long chain polyunsaturated FAs (4 out of 4 and 6 out of 8 significant) and long chain monounsaturated FAs (6 out of 6). After the higher exposure, unsaturated CARs were either significantly downregulated (4 out of 10) or trended downwards (6 out of 10) compared to the sham-exposure. The accumulation of FAs in Etomoxir-treated mice after 112 dB noise indicates that the reason lipid changes were not seen in untreated animals at this exposure was due to increased mobilization of FAs rather than a lack of β-oxidation. Together, these data indicate that β-oxidation is utilized during both moderate and intense noise, with FA mobilization increasing with noise intensity (Figure 3A-C).

**Figure 3:**
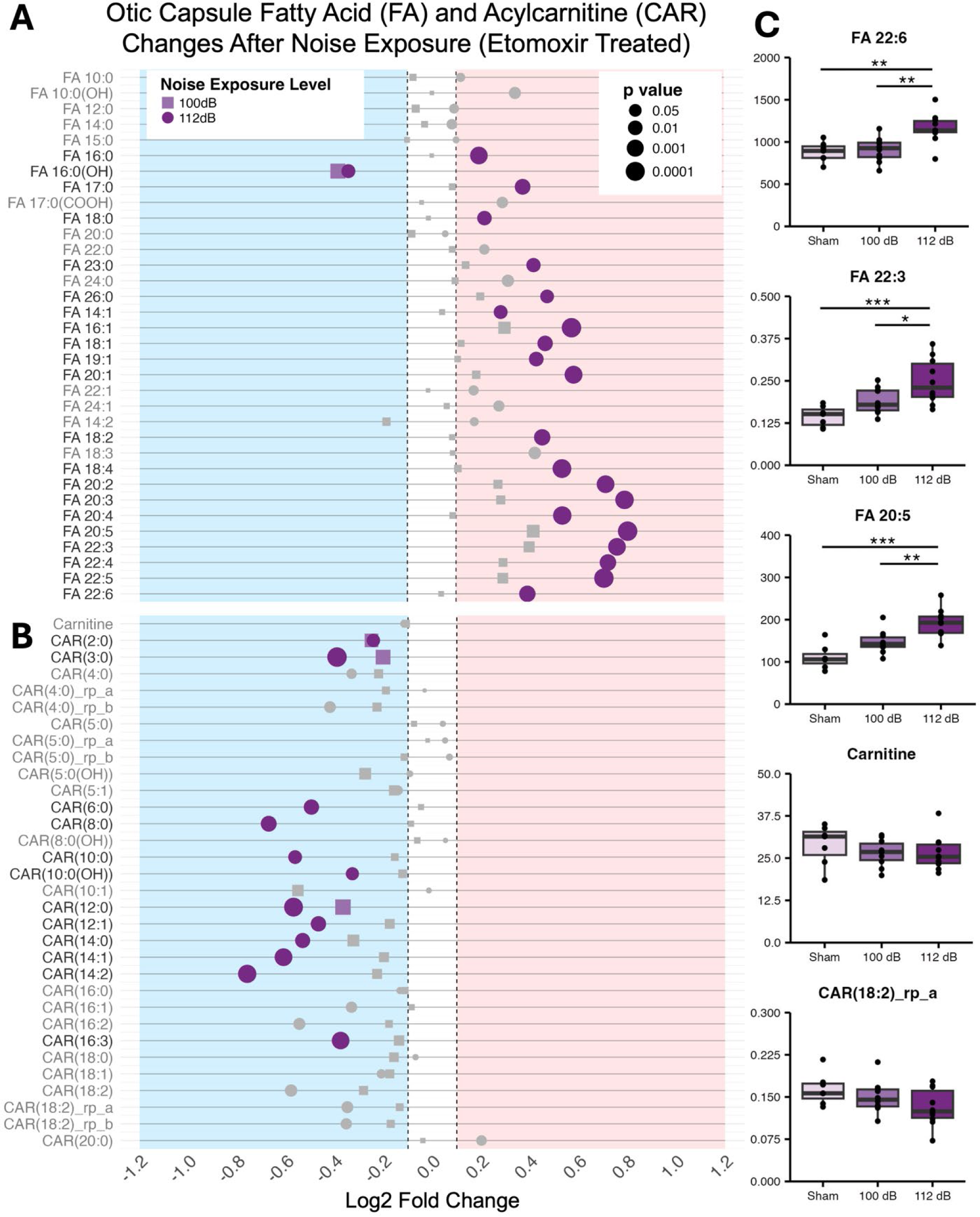
Noise exposure in mice unable to utilize β-oxidation (Etomoxir treatment) increased FA and decreased CAR concentrations, supporting the hypothesis that noise increases otic capsule β-oxidation. **(A-B)** Noise-induced fatty acid (FA) and acylcarnitine (CAR) changes are eliminated or reversed in Etomoxir-treated mice, indicating that noise activates beta-oxidation. Sham-exposed (Etomoxir-treated) animals were used as controls. Two-way ANOVA with Tukey’s multiple comparison test was used for statistical analysis. Dots represent lipid species and are color coded when adjusted-p<0.05. The dot’s shape and color correspond to the noise exposure level. **(C)** Quantitative levels of selected lipids in noise- and sham-exposed otic capsules. Values are in µg (lipid) / mg (sample). Two-way ANOVA with Tukey’s multiple comparison test was used for statistical analysis. Dots represent otic capsule samples. *p < 0.05; **p < 0.01; ***p < 0.001.

### Chronic Etomoxir treatment does not affect baseline hearing but reduces the degree of NIHL caused by a 112 dB SPL exposure

Β-oxidation is an important source of cellular energy, but it produces reactive oxygen species (ROS) that can be detrimental to cellular and tissue function^22,23^. To determine the impact of loss of β-oxidation on hearing and NIHL, we compared the auditory function of mice treated with Etomoxir or saline under normal acoustic conditions (no noise exposure). Specifically, auditory brainstem responses (ABRs) and distortion product otoacoustic emissions (DPOAEs) were recorded. ABRs reflect the function of the ascending auditory pathway, from the activation of IHCs and SGNs to the inferior colliculus, whereas DPOAEs reflect OHCs function^2^.

To evaluate the impact of Etomoxir on hearing under normal acoustic conditions, mice were treated with Etomoxir or saline daily for 14 days, and ABR and DPOAE recordings were taken at baseline and on the final day of treatment. Etomoxir treatment did not affect ABR and DPOAE thresholds, nor the amplitudes of the first peak of the ABR waveform, which reflect the intensity of the synchronous activity of SGNs in response to sound (Supplemental Figure 2). These results demonstrate that Etomoxir has no effect on hearing per se and support the hypothesis that β- oxidation is not critical under normal acoustic conditions.

Next, we evaluated the effects of chronic Etomoxir on the effects of 112 dB SPL noise exposure, which causes a small degree of hair cell damage and permanent elevation of auditory thresholds^4^. Etomoxir or saline (vehicle) treatment was continued for 2 more days (16 days total) before exposure to noise (112 dB SPL; 8-16 kHz; 2 hours). ABR and DPOAE recordings were taken before noise exposure as baseline, and at 1, 7, and 14 days after exposure. Surprisingly, Etomoxir treatment provided significant protection from 112 dB SPL noise exposure (Figure 4A). Specifically, ABR and DPOAE thresholds shifts were smaller in the treated animals at all timepoints throughout the 14-day recovery period, with a maximal protective effect of ∼20 dB, reflecting a significant improvement in hearing compared to mice treated with vehicle. These results suggest that noise-induced β-oxidation in the inner ear enhances the degree of NIHL caused by this type of exposure.

**Figure 4:**
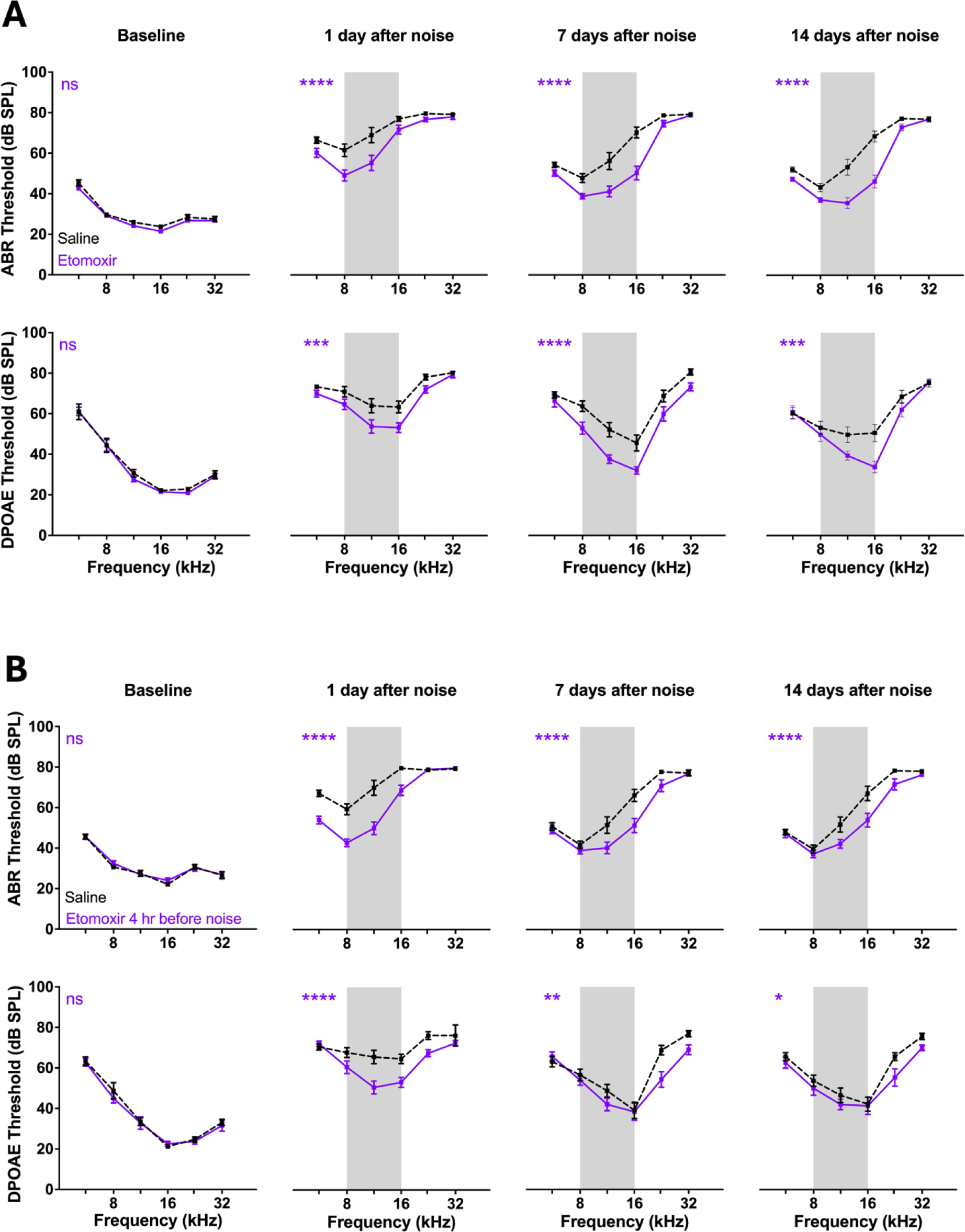
Etomoxir treatment reduces the degree of hearing loss induced by a 2-hour exposure to 112 dB SPL, 8-16 kHz. Chronic prophylactic Etomoxir treatment reduces the degree of noise-induced hearing loss. Mice were treated daily with Etomoxir (35 mg/kg IP) or saline (vehicle) for 16 days before noise exposure (112 dB SPL; 8-16 kHz (grey band); 2 hours). ABR and DPOAEs were recorded on the last day of treatment (baseline), and days 1, 7, and 14 after noise. Higher thresholds represent reduced hearing ability. Etomoxir, n=17 mice; Saline, n=16 mice. **(B)** One dose of Etomoxir is sufficient to protect the inner ear against noise-induced damage. Mice were treated with Etomoxir (35 mg/kg IP) or saline (vehicle) 4 hours before noise exposure (112 dB SPL; 8-16 kHz, grey band, 2 hours). ABR and DPOAEs were recorded before treatment (baseline), and on days 1, 7, and 14 after noise. Etomoxir, n=15 mice; Saline, n=16 mice. Data are shown as mean ± SEM and were evaluated at each time point by mixed model 2-way ANOVA followed by Šídák multiple- comparison correction. ns = not significant; *p < 0.05; **p < 0.01; ***p < 0.001; ****p < 0.0001.

### Etomoxir treatment reduces the degree of NIHL when injected acutely before but not after noise exposure

When considering approaches to protect the inner ear from anticipated noise exposure, a drug that works with a single prophylactic dose, or with treatment after exposure, would be preferable to one that requires chronic prophylactic treatments. Therefore, we evaluated the effectiveness of a single Etomoxir injection (35 mg/kg; IP) either 4 hours before or 24 hours after exposure to 112 dB SPL (8-16kHz; 2 hours). Remarkably, a single dose of Etomoxir before the exposure was sufficient to significantly protect both ABR and DPOAE thresholds (Figure 4B), though to a lesser degree than chronic treatment (Figure 4A). In contrast, CPT-1A inhibition after noise exposure did not affect the noise-induced threshold shifts (Supplemental Figure 3).

### Pre-exposure Etomoxir treatment reduces the extent of noise-induced hair cell loss

To begin to explore the potential effect of prophylactic Etomoxir treatment on ear structure, we examined the impact of noise exposure on the number of inner and outer hair cells in mice treated 4 hours before the exposure with saline or Etomoxir. Otic capsules were harvested 1 week after the last recording (3 weeks after noise exposure), the cochleae were microdissected, immunostained for the hair cell marker Myosin VIIa, and the number of IHCs and OHCs quantified using confocal microscopy. Organ of Corti from Etomoxir-treated mice had reduced IHC loss in the 32 kHz region of the cochlea, and there was a trend towards protecting HCs at the other frequencies, indicating that Etomoxir can improve hair cell survival after noise exposure (Supplemental Figure 4).

## Discussion

The prevalence of NIHL in both young and older individuals, together with the lack of regenerative capacity in the organ of Corti, makes finding ways to prevent or reduce the damaging effects of noise on the inner ear an urgency. Using untargeted and targeted lipidomics, we demonstrate that noise exposure increases β-oxidation in the inner ear. Furthermore, we show that inhibition of β- oxidation by Etomoxir reduces the degree of NIHL, indicating that lipid metabolism plays a role in the pathogenesis of this highly prevalent condition.

While β-oxidation of long-chain FAs serves as an energy source, our results indicate that this oxidative cycle contributes to NIHL. β-oxidation is known to generate reactive oxygen species (ROS) that can cause cell damage and death^22,23^. ROS-related pathways have been repeatedly identified as contributing to hearing loss associated with aging and noise, typically by inducing HC dysfunction and loss^5,24–28^, though the metabolic sources of ROS in NIHL remain unclear. The protective effect of inhibiting β-oxidation suggests that this pathway may significantly contribute to ROS production during noise exposure. Furthermore, the lack of reported hearing deficits in patients unable to utilize long, medium, and short-chain β-oxidation^29,30^ suggests that inhibition of β-oxidation may be an effective strategy to reduce ROS and alleviate NIHL in humans without altering baseline hearing. This is also in line with the observation that knockout of fatty acid binding protein 7 (FABP7), which reduces polyunsaturated fatty acid transport, mitigates cochlear damage after noise and enhances HC survival in mice^31^. Additionally, metabolic dysfunction leading to impaired FA oxidation, accumulation of FAs, and dysregulated gluconeogenesis has also been linked to increased HC loss in age-related hearing loss models^32^.

The energetic requirements of the inner ear during noise exposure, and the molecular mechanisms by which it meets them, remain poorly understood. Particularly, the role of lipids and β-oxidation in NIHL remained a substantial knowledge gap, despite several studies suggesting these pathways may play an important role. Namely, studies of guinea pigs perilymph reported noise-induced increases in 3-hydroxy-butyrate^33^, in several short-chain CARs^34^, and in glycerophospholipid metabolism^35^. Moreover, acetoacetate has been reported to be a viable fuel source for OHCs *in vivo*^21^, and several studies tested if acetyl-l-carnitine treatment reduces NIHL, both alone and in combination with other antioxidants, some observing a protective effect when administered alone^36–40^. Furthermore, even though it is well established that noise increases metabolic demands, there is an unexplained biphasic relationship between glucose uptake and noise exposure intensity in mice. Specifically, previous studies reported that glucose uptake increases with intensity until a certain exposure level is reached (∼70 dB SPL), at which point increasing intensity attenuates uptake^15^. Our results showing β-oxidation is increased during noise exposure could be related to the reduced glucose metabolism. The noise-induced increased FA uptake could meet some of the energetic demands caused by reduced glucose availability under the aegis of noise stress at the expense of generation of excessive ROS and damage to the particularly sensitive cells^21,25,41–46^. We see that the protection provided by Etomoxir treatment on the cochlea is greatest for IHCs and their function, suggesting that IHCs are more sensitive to the noise-induced effects of β-oxidation than OHCs. In contrast, animal studies reported antioxidants as having stronger protective effects on OHCs^24,25,44^. Antioxidants have not been successful in preventing NIHL in human trials^47^. Thus, it would be interesting to evaluate if a combination of Etomoxir (to prevent long-chain β-oxidation) and antioxidants (to “clean up” remaining ROS) provides greater protection than each independently. Furthermore, since the protective effects of Etomoxir on auditory thresholds are larger than the effects on HC survival, our results suggest that β-oxidation may primarily cause non-apoptotic damage, such as to HC stereocilia, or by affecting HC metabolic function. Further ultrastructural and metabolic interrogations of Etomoxir-treated inner ears may uncover the mechanism by which noise-induced β-oxidation damages the cochlea.

Though Etomoxir is considered to be safe and well tolerated in animals, its usage was discontinued in phase 2 human clinical trials due to hepatotoxicity^20^. Our findings that the FA changes occur locally in the otic capsule tissue raise the possibility that Etomoxir could be administered locally, e.g., via ear drops, which would reduce the possibility of systemic effects.

In summary, our results identifying β-oxidation as a source of energy during noise stress and the protective effect of its inhibition with β-oxidation via Etomoxir demonstrate that lipidomics profiling can be a powerful tool to understand and intervene on inner ear pathology. As there are no FDA approved pharmacological therapies to prevent, reduce, or treat NIHL, our identification of potential therapeutic strategies represents an important future direction of interrogation.

## Materials and Methods

### Experimental groups and noise exposure

All animal procedures were approved by the Institutional Animal Care and Use Committee of the University of Michigan, and all experiments were performed in accordance with relevant guidelines and regulations. We used female CBA/J mice with normal hearing housed in a facility with ambient sound levels of 40–50 dB SPL within frequencies detectable by the mouse ear. Animals were randomly assigned to a control or noise-exposed group, or to a treatment (Etomoxir) or vehicle (saline) group.

Awake mice were placed within small cells in a subdivided cage, suspended in a reverberant noise exposure chamber. Noise exposure varied from 98 to 112 dB SPL, with sham-exposed groups being placed in the chamber with the speaker off. Noise or sham duration was always 2 hours, and noise exposures were to octave-band noise (8–16 kHz). Noise calibration to target SPL was performed before each exposure session. Sound pressure levels varied by ∼1 dB SPL across the cages.

Experimental groups and corresponding data are summarized below. Animals in experimental groups interrogated simultaneously are indicated by numbered group. Etomoxir (Cayman, item 11969) was reconstituted at a concentration of 2.5 mg/mL in 0.9% saline and always administered at a dose of 35 mg/kg, with volume-matched saline controls. Treatment occurred before noise exposure unless otherwise indicated.

**Table 1:**
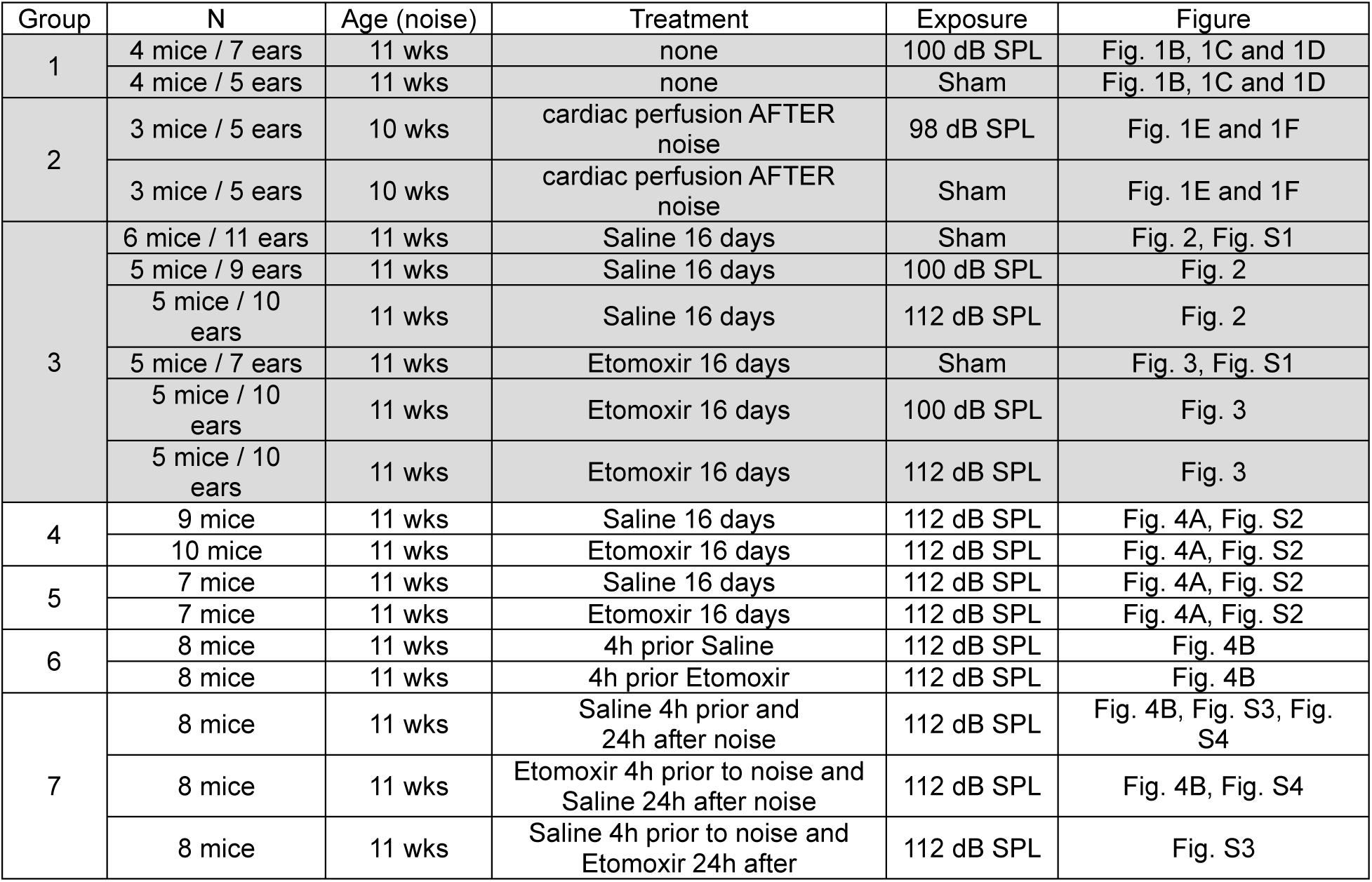
Experimental groups, animal numbers, treatment, and noise-exposure conditions. Summary of the experimental groups, number of mice, and treatment and noise-exposure conditions. Shared group numbers indicate the animals were interrogated simultaneously. Treatments occurred before noise exposure unless otherwise indicated. All noise or sham exposures were performed for 2 hours. Shaded rows reflect animals used for profiling, and white rows reflect animals used for physiological testing.

| Group | N | Age (noise) | Treatment | Exposure | Figure |
| --- | --- | --- | --- | --- | --- |
| 1 | 4 mice / 7 ears | 11 wks | none | 100 dB SPL | Fig. 1B, 1C and 1D |
|  | 4 mice / 5 ears | 11 wks | none | Sham | Fig. 1B, 1C and 1D |
| 2 | 3 mice / 5 ears | 10 wks | cardiac perfusion AFTER noise | 98 dB SPL | Fig. 1E and 1F |
|  | 3 mice / 5 ears | 10 wks | cardiac perfusion AFTER noise | Sham | Fig. 1E and 1F |
| 3 | 6 mice / 11 ears | 11 wks | Saline 16 days | Sham | Fig. 2, Fig. S1 |
|  | 5 mice / 9 ears | 11 wks | Saline 16 days | 100 dB SPL | Fig. 2 |
|  | 5 mice / 10 ears | 11 wks | Saline 16 days | 112 dB SPL | Fig. 2 |
|  | 5 mice / 7 ears | 11 wks | Etomoxir 16 days | Sham | Fig. 3, Fig. S1 |
|  | 5 mice / 10 ears | 11 wks | Etomoxir 16 days | 100 dB SPL | Fig. 3 |
|  | 5 mice / 10 ears | 11 wks | Etomoxir 16 days | 112 dB SPL | Fig. 3 |
| 4 | 9 mice | 11 wks | Saline 16 days | 112 dB SPL | Fig. 4A, Fig. S2 |
|  | 10 mice | 11 wks | Etomoxir 16 days | 112 dB SPL | Fig. 4A, Fig. S2 |
| 5 | 7 mice | 11 wks | Saline 16 days | 112 dB SPL | Fig. 4A, Fig. S2 |
|  | 7 mice | 11 wks | Etomoxir 16 days | 112 dB SPL | Fig. 4A, Fig. S2 |
| 6 | 8 mice | 11 wks | 4h prior Saline | 112 dB SPL | Fig. 4B |
|  | 8 mice | 11 wks | 4h prior Etomoxir | 112 dB SPL | Fig. 4B |
| 7 | 8 mice | 11 wks | Saline 4h prior and 24h after noise | 112 dB SPL | Fig. 4B, Fig. S3, Fig. S4 |
|  | 8 mice | 11 wks | Etomoxir 4h prior to noise and Saline 24h after noise | 112 dB SPL | Fig. 4B, Fig. S4 |
|  | 8 mice | 11 wks | Saline 4h prior to noise and Etomoxir 24h after | 112 dB SPL | Fig. S3 |

### Lipidomics: Inner ear extraction

Temporal bones were harvested immediately after the exposure and the auditory bulla were isolated from the surrounding skull bone and placed in a Petri dish with phosphate-buffered saline (PBS). We then quickly removed any attached musculature or soft tissue, the middle ear and the cerebellar parafloculli to isolate the bony otic capsule containing the entire cochlea and vestibular organs (including sensory epithelia, sensory neurons, nerves, etc.) for processing. The round and oval windows were kept intact during the dissection to prevent dilution or contamination of the inner ear fluid with the bath solution. A single inner ear tissue from a mouse was placed in an individual tube and flash frozen. Mice were not cardiac perfused after noise exposure except where indicated (mice in group 2a and 2b).

### Targeted Lipidomics: Sample preparation and extraction

Otic capsules are carefully weighed in a microtube containing stainless steel beads for tissue disruption, and extraction solvent (8:2 methanol/water containing isotope-labeled internal standards) is added in a ratio of 30 μl/mg. Tissue disruption is performed in a Precellys Evolution Touch homogenizer set to 0°C, 6200 rpm, for 3 cycles (20 sec shake time and 30 sec rest time per cycle.) After removal from the homogenizer, samples are incubated on ice for 10 minutes and vortexed to remix, then finally centrifuged at 14,000 RPM for 10 min at 4°C. 200 μL of extract is transferred to an autovial and dried under a continuous stream of nitrogen at room temperature for 2 hours. Dried samples are reconstituted in 100 μL of 80:20 water/methanol.

A pooled sample for quality control is created by combining 20 μL of extract from each sample and processed alongside the other samples. This sample was run periodically throughout the mass spectrometry analysis.

### Targeted Lipidomics: Optimized LC-MS methods

LC-MS analysis was performed on an Agilent system consisting of an Agilent 1290 Infinity II / 6545 qTOF MS system with the JetStream Ionization (ESI) source (Agilent Technologies, Inc., Santa Clara, CA USA) using an Acquity HSS T3, 2.1 mm x 100 mm, 1.7 µm particle size column (Waters Corporation, Milford, MA). Each sample was analyzed twice, once in positive and once in negative ion mode. Mobile phase A was 100% water, B was 100% methanol, and C was 2.5% formic acid. The gradient for both positive and negative ion modes was as follows: .0% B, 4% C (0 min), 99% B, 1C% (10 min), 99%B, 1%C (17 min), 0%B, 1%C (17.1min), The column was then reconditioned for 3 min at starting conditions before moving to the next injection. The flow rate was 0.45 mL/min and the column temperature was 55°C. The injection volume for positive and negative modes was 10 µL. Source parameters were drying gas temperature 275°C, drying gas flow rate 12 L/min, nebulizer pressure 45 psig, sheath gas temp 325°C and flow 12 l/min, and capillary voltage 4000V, with internal reference mass correction.

### Targeted Lipidomics: Internal Standards

For Fatty Acid analysis, a custom mix was made in house containing D5-DHA, D5-EPA, D8- Archidonic acid, D31-palmitic acid, D35-stearic acid, D27-Myristric acid in with concentration in the extraction solvent ranging from 160 to 600 nM.

For acylcarnitine analysis, Cambridge Isotopes NSK-B1 mix was used (concentrations ranging from 7.6 to 152 µM, depending on the metabolite) and was added to the extraction solvent at a ratio of 6.25 μL /30 mL, resulting in metabolite concentrations ranging from 1.6 to 31 nM in the extraction solvent.

### Targeted Lipidomics: Metabolite Quantification

Metabolite areas for both the endogenous compounds and isotopically labeled internal standards were obtained by manual integration using Profinder v8.00 (Agilent Technologies, Santa Clara, CA.) Metabolites were identified by matching the retention time (+/- 0.1 min), mass (+/- 10 ppm) and isotope profile (peak height and spacing) to authentic standards. Metabolite quantitation was obtained by 1-point calibration to the nearest internal standard.

### Untargeted Lipidomics: Sample preparation and extraction

Otic capsules were accurately weighed and then homogenized. Lipids were extracted using a modified Bligh-Dyer method^48^ using a 2:2:2 ratio volume of methanol/water/dichloromethane at room temperature after spiking internal standards (described above) The organic layer was collected and completely dried under nitrogen. Before MS analysis, the dried lipid extract was reconstituted in 100 mL of buffer B (10:85:5 acetonitrile/ isopropyl alcohol/water) containing 10mMammonium acetate and subjected to LC-MS.

### Untargeted Lipidomics: Data Dependent Liquid chromatography-mass spectrometry (LC-MS/MS) for measurements of lipids

Chromatographic separation was performed on a Shimadzu CTO-20A Nexera X2 UHPLC system equipped with a degasser, binary pump, thermostat autosampler, and column oven [all components manufactured by Shimadzu (Canby,OR)]. The column heater temperature was maintained at 55°C and an injection volume of 5 mL was used for all analyses. For lipid separation, the lipid extract was injected onto a 1.8-mm particle diameter, 50-mm 3 2.1-mm inner diameter Waters Acquity HSS T3 column (Waters, Milford, MA). Elution was performed using acetonitrile/water (40:60, v/v) with 10 mM ammonium acetate as solvent A and acetonitrile/water/isopropanol alcohol (10:5:85, v/v/v) with 10 mM ammonium acetate as solvent B. For chromatographic elution we used a linear gradient during a 20-minute total run time, with a 60% solvent A and 40% solvent B gradient in the first 10 minutes. Then the gradient was ramped in a linear fashion to 100% solvent B, which was maintained for 7 minutes. Thereafter, the system was switched back to 60% solvent B and 40% solvent A for 3 minutes. The flow rate used for these experiments was 0.4 mL/min and the injection volume was 5 mL. The column was equilibrated for 3 minutes before the next injection and run at a flow rate of 0.400 mL/min for a total run time of 20 minutes. MS data acquisition for each sample was performed in both positive and negative ionization modes using a TripleTOF 5600 equipped with a DuoSpray ion source (AB Sciex, Concord, ON, Canada). Column effluent was directed to the electrospray ionization source and voltage was set to 5500 V for positive ionization and 4500 V for the negative ionization mode. The declustering potential was 60 V and the source temperature was 450°C for both modes. The curtain gas flow, nebulizer, and heater gas were set to 30, 40, and 45, respectively (arbitrary units). The instrument was set to perform one time of flight MS survey scan (150 ms) and 15 MS/MS scans with a total duty cycle time of 2.4 seconds. The mass range of both modes was 50 to 1200 m/z. Acquisition of MS/MS spectra was controlled by the data-dependent acquisition function of the Analyst TF software (AB Sciex) with application of the following parameters: dynamic background subtraction, charge monitoring to exclude multiply charged ions and isotopes, and dynamic exclusion of former target ions for 9 seconds. Collision energy spread of 20 V was set whereby the software calculated the collision energy value to be applied as a function of m/z. A DuoSpray source coupled with automated calibration system (AB Sciex) was used to maintain mass accuracy during data acquisition. Calibrations were performed at the initiation of each new batch or polarity change.

### Untargeted Lipidomics: Internal Standards

Quality control samples were prepared by pooling equal volumes of each sample and injected at the beginning and the end of each analysis and after every 10 sample injections to provide a measurement of the system’s stability and performance as well as reproducibility of the sample preparation method^49^. Two kinds of controls were used to monitor the sample preparation and MS. To monitor instrument performance, 10 mL of a dried matrix-free mixture of the internal standards reconstituted in 100 mL of buffer B (85% isopropyl alcohol/10% acetonitrile/5% water in 10mMNH4OAc) was analyzed. As additional controls to monitor the profiling process, an equimolar mixture of 13 authentic internal standards and a characterized pool of human plasma and test pool (a small aliquot from the all-adipose tissue samples used in this study; extracted in tandem with tissue samples) were analyzed along with adipose tissue samples. Each of these controls was included several times into the randomization scheme such that sample preparation and analytical variability could be monitored constantly.

### Untargeted Lipidomics: Normalization

Each lipid was normalized using an Internal standard that minimized its RSD post-normalization. Each mode data was normalized separately. Normalized data from both modes is combined, then repeats removed to give the final dataset.

### Untargeted Lipidomics: Data Cleaning and Degeneracy Removal

First step in data analysis is filtering features based on missing values. There are two types of QC samples run along with the experimental samples. Sample ID’s CS00000001 are called “Test.Pool” samples and CS00000004 are called “Pooled.Plasma”.“Pooled.Plasma” is a red cross plasma pool run as an internal QC sample in all studies. “Test.Pool” is made by taking small aliquot from each experimental sample and pooling it together to create a QC sample which is run after every few runs to get assess drifts and other variation caused by the run. Features missing in more than 30% Test.Pool samples are removed. Missing patterns are carefully examined for groupwise missingness. The data is imputed using KNN algorithm with knn.impute function from bnstruct package.

### Untargeted Lipidomics: Lipid identification

The raw data were converted to mgf data format using ProteoWizard software^50^. The NIST MS PepSearch program was used to search the converted files against LipidBlast^51,52^ libraries in batchmode. We optimized the search parameters using the NIST11 library and LipidBlast libraries and comparing them against our lipid standards. To facilitate accurate lipid identification, a stringent mass error tolerance of 0.001 m/z for positive mode and 0.005 m/z for negative mode were used. The minimum match factor used in the PepSearch Program was set to 200. The MS/MS identification results from all of the files were combined using an in-house software tool to create a library for quantification. All raw data files were searched against this library of identified lipids with mass and retention time using MultiQuant 1.1.0.26 (AB Sciex)^53^. Quantification was done using MS1 data. The quality control samples were also used to remove technical outliers and lipid species that were detected below the lipid class–based lower limit of quantification. Quality control samples evenly distributed along analytical runs of the study were analyzed. The average coefficient of variation of all the lipids detected in the study samples was 25%.

### Untargeted Lipidomics: Bioinformatic and statistical analysis

Quality control was performed by removing lipid species with >40% relative standard deviation in pooled QC samples. Lipid intensities were normalized to the mean total ion current across all samples.

For individual lipids, significance was assessed by two-tailed t-test without multiple testing corrections.

For grouped analysis: lipids were assigned to *a priori* lipid groups based on headgroup, chain length, and saturation as below. For each group, lipid intensities were summed within each sample to generate group values, which were then analyzed by two-tailed t-test without multiple testing corrections.

Headgroups: PC, phosphatidylcholine; PE, phosphatidylethanolamine; PI, phosphatidylinositol; PG, phosphatidylglycerol; LPC, lysophosphatidylcholine; LPE, lysophosphatidylethanolamine; LNAPE, N-acyl phosphatidylethanolamine; O-, ether; Cer, ceramide; CerP, ceramide phosphate; SM, sphingomyelin; DG, diacylglycerol; TG, triacylglycerol; FA, fatty acid.

Chain Length: Very Long: >20 carbons (C)/side chain (average); Long: 12-20, Medium: 6-12.

Saturation: Polyunsaturated: >1 units unsaturation/chain (average), Monounsaturated: 0-1, Saturated: 0.

All analyses were performed using R software, version 3.5.3 (The R Foundation for Statistical Computing) unless otherwise noted. Significance was defined as p<0.05.

### Immunostaining and hair cell counts

Inner ear tissues were dissected as above and fixed in 4% paraformaldehyde (instead of flash frozen) in 0.1 M phosphate buffer for 2 h at room temperature, followed by decalcification in 5% EDTA at 4 °C for 5 days. Cochlear tissues were microdissected and permeabilized by freeze– thawing in 30% sucrose. The microdissected tissues were incubated in blocking buffer containing 5% normal horse serum and 0.3% Triton X-100 in PBS for 1 h. Tissues were then incubated with anti-myosin VIIa (rabbit anti-MyoVIIa; Proteus Biosciences, Ramona, CA; 1:500) primary antibodies (diluted in blocking buffer) at 37 °C for 16 h.

Tissues were then incubated with appropriate Alexa Fluor-conjugated fluorescent secondary antibody (Invitrogen, Carlsbad, CA; 1:1000 diluted in 1% normal horse serum and 0.3% Triton X- 100 in PBS; AF647 IgG catalog no. A-212z4) for 1 h at room temperature. The tissues were mounted on microscope slides in ProLong Diamond Antifade Mountant (Thermo Fisher Scientific). All pieces of each cochlea were imaged at low power (10X magnification) to convert cochlear locations into frequency (tonotopic mapping) using a custom plug-in to ImageJ (1.53c NIH, MD) available at the website of the Eaton-Peabody Laboratories (EPL). Cochlear tissues from the 5.6, 8.0, 11.3, 16.0, 22.6, and 32.0 regions, corresponding to the ABR/DPOAE recordings, were used for further analyses.

Confocal z-stacks (0.3 μm step size) of cochlear tissues were taken using a Leica SP8 microscope equipped. Images for quantitative analyses of the nodal structures were taken under × 20 lens with × 2 optical zoom. ImageJ software (version 1.54g, NIH, MD) was used for image processing and three-dimensional reconstruction of z-stacks.

### Distortion product otoacoustic emissions (DPOAEs) and auditory brainstem responses (ABRs)

DPOAEs and ABRs were performed as previously described ^54^. Mice were anaesthetized by i.p. injections of xylazine (20 mg kg−1, i.p.) and ketamine (100 mg kg−1, i.p.).

For ABR recordings, subdermal electrodes were placed (1) at the dorsal midline of the head; (2) behind the left earlobe; and (3) at the base of the tail (for a ground electrode). ABRs were evoked with 5 ms tone pips (0.5 ms rise–fall) delivered to the eardrum through a closed acoustic system, consisting of two sound sources (CDMG15008-03A, CUI) and an electret condenser microphone (FG-23329-PO7, Knowles) as an in-dwelling probe microphone. The frequencies of tone pips were 5.6, 8, 11.3, 16, 22.6, and 32, with sound levels from 20 to 80 dB SPL for each frequency.

The response was amplified (10,000X) and filtered (0.3–3 kHz) with an analog-to-digital board in a PC-based data-acquisition system. The sound level was raised in 5 dB step intervals from 20- 80 dB SPL. At each level, the average of 1,024 signals was taken after “artifact rejection.” ABR threshold was determined by visual analysis of stacked waveforms from highest to lowest SPL. Threshold was determined as the lowest level at which a repeatable Wave I could be identified. Wave I–V amplitude was defined as the difference between a 1-ms average of the pre-stimulus baseline and the wave I–V peak, after additional high-pass filtering to remove low-frequency baseline shifts.

For DPOAEs, otoacoustic emissions in response to 2 primary tones (f1 and f2) and recorded at (2 × f1)−f2. f1 level was 10 dB higher than the f2 level and frequency ratio f2/f1 was 1.2. The ear-canal sound pressure was amplified and averaged at 4 μs intervals. DPOAE thresholds were defined as the f2 level that produced a response consistently 3 dB SPL higher than the noise floor.

Both ABR and DPOAE recordings were performed using the EPL cochlear function test suite (Mass Eye and Ear, Boston, MA, USA). ABR thresholds and ABR peak I amplitudes were analyzed with ABR peak Analysis software (Mass Eye and Ear, Boston, MA, USA). DPOAE thresholds were analyzed using custom MatLab script. Statistical analyses were performed using GraphPad Prism version 10.4.1.

## Author Contributions

GW, LJ, and GC designed the research studies with input from CFB and CAL. GW, LRJ, and LC conducted experiments. MB performed the profiling. GW performed raw data analysis. GW, CFB, and GC performed data analysis and interpretation. GW, LJ, LRC, MB, CAL, CFB, and GC wrote the manuscript.

### Acknowledgments

This work was supported by NIH R01DC000188 (GC) and R21DC017916 (GC).

---

**Supplemental Figure 1:**
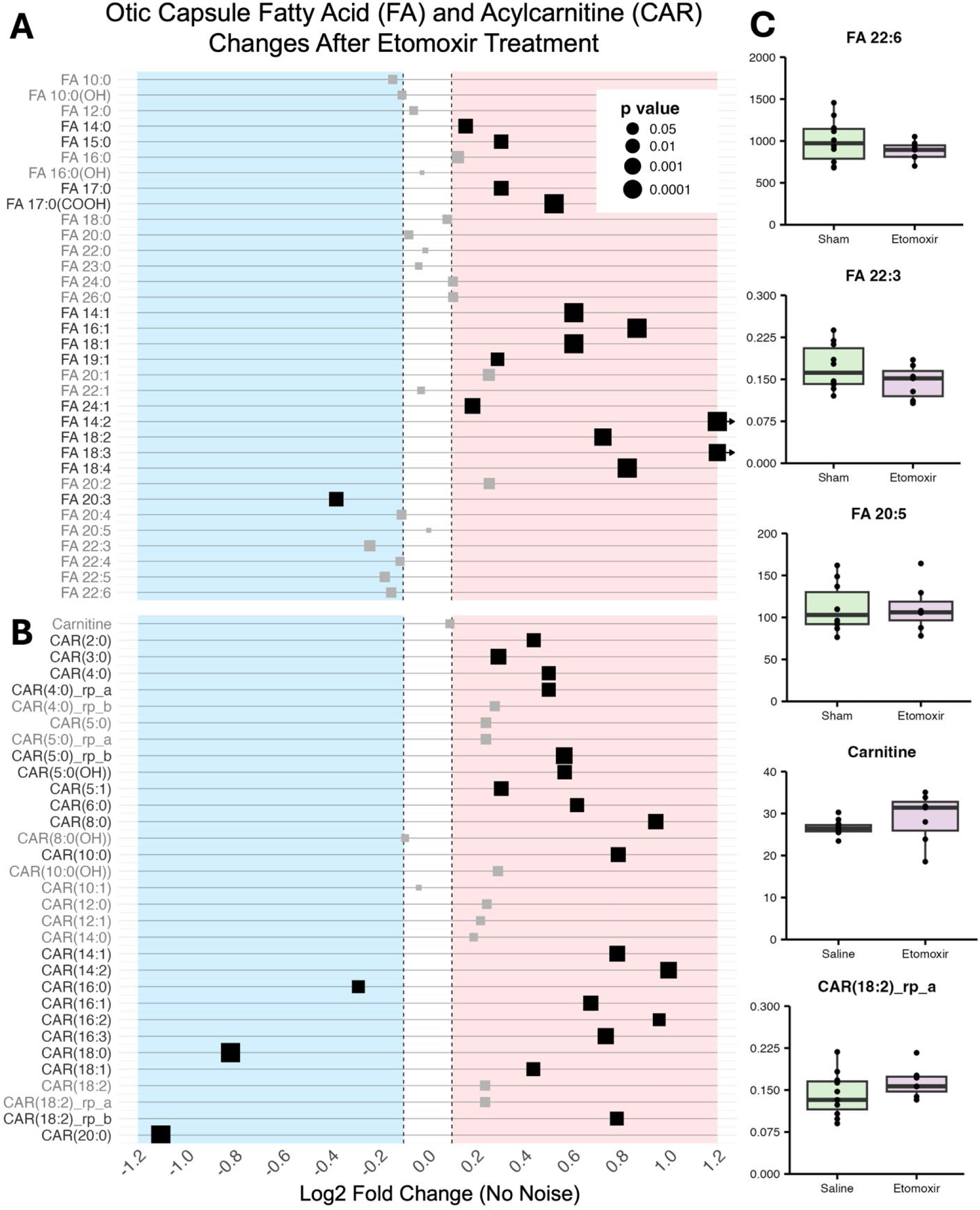
Under normal acoustic conditions, Etomoxir does not alter the baseline concentration of very long polyunsaturated fatty acids, and inconsistent FA and CAR changes suggest a lack of β-oxidation. **(A-B)** Fatty acid (FA) and acylcarnitine (CAR) species change after treatment with Etomoxir to inhibit CPT-1A and prevent long-chain FA beta-oxidation. Saline-treated (vehicle) animals were used as controls. Student’s t-test with Benjamini-Hockberg correction was used for statistical analysis. Each dot represents one lipid species and is color coded when adjusted-p<0.05. **(C)** Quantitative levels of selected lipids after Etomoxir treatment. Values are in µg (lipid) / mg (sample). Student’s t-test with Benjamini-Hockberg correction was used for statistical analysis. Each dot represents one otic capsule sample.

**Supplemental Figure 2:**
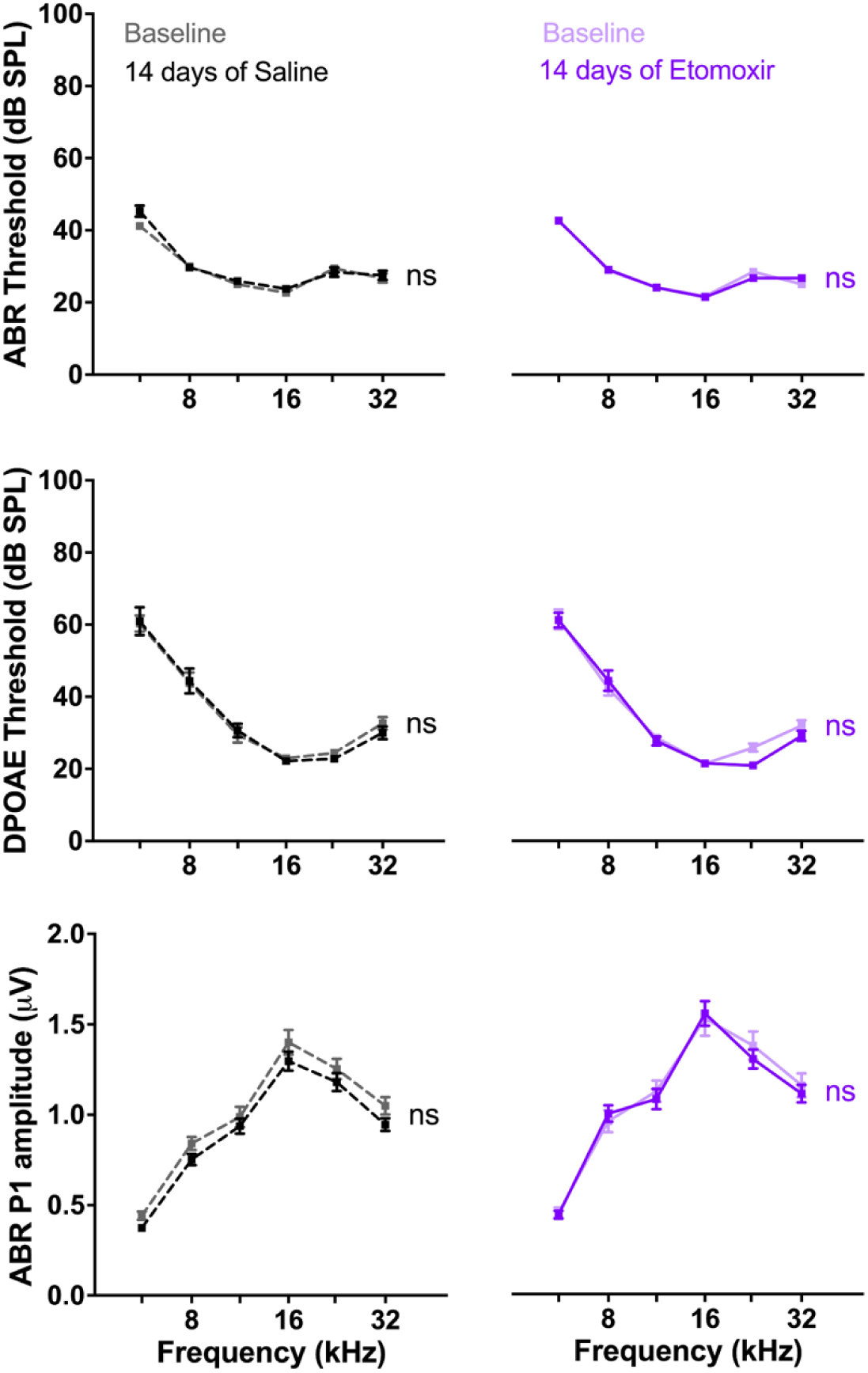
Etomoxir treatment, which prevents utilization of long-chain FA beta-oxidation via inhibition of CPT-1A, does not alter hearing. Mice were treated daily with Etomoxir (35 mg/kg IP) or saline (vehicle) for 14 days. ABR and DPOAEs were recorded before treatment (baseline) and on day 14 of treatment. Data are shown as mean ± SEM and were evaluated for each treatment group by mixed model 2-way ANOVA followed by Šídák multiple-comparison correction. Etomoxir, n=17 mice; Saline, n=16 mice. ns = not significant.

**Supplemental Figure 3:**
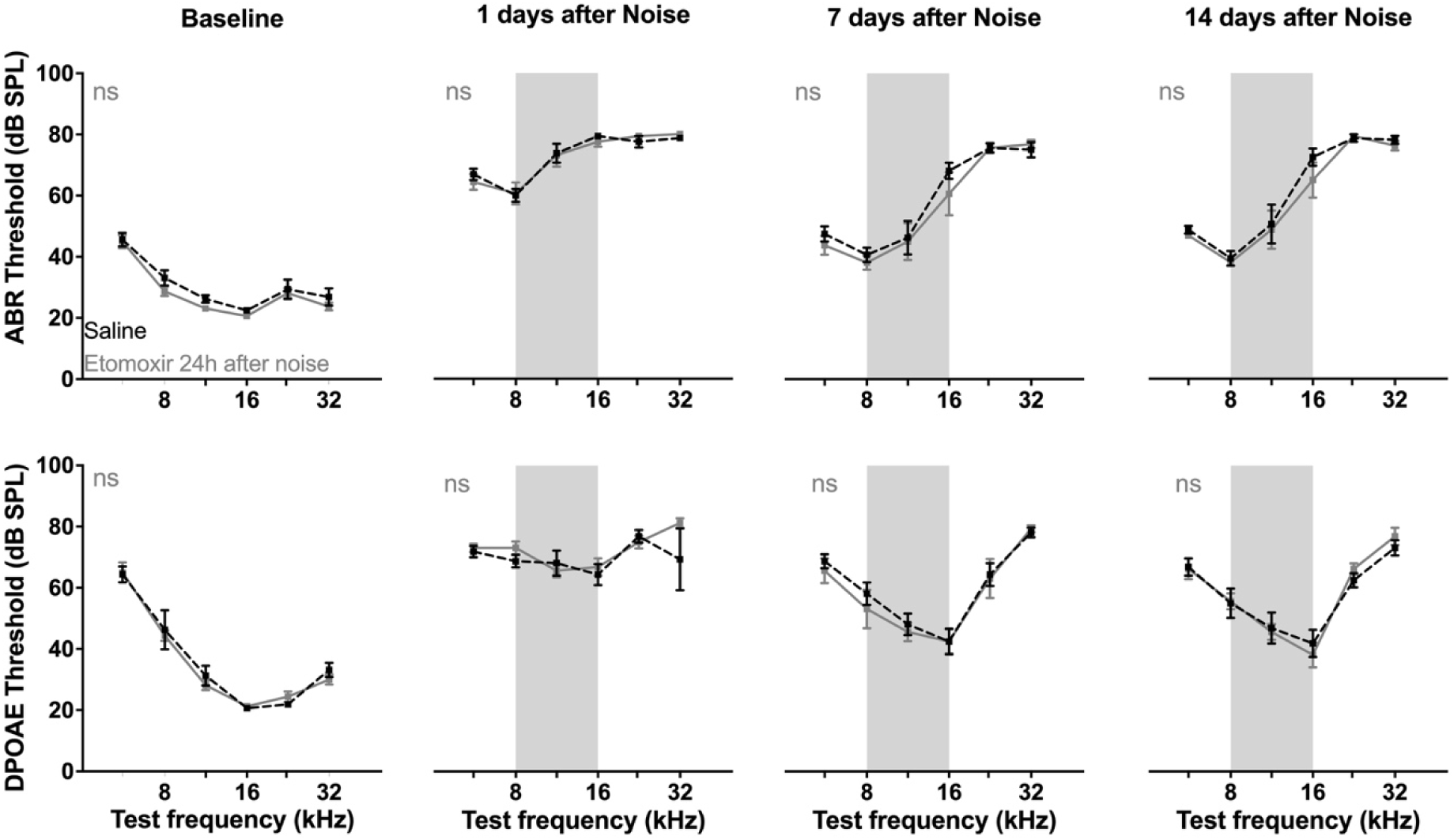
Etomoxir treatment 24 hours after noise does not reduce the extent of hearing loss induced by a 2-hour exposure to 112 dB SPL, 8-16 kHz. Post-treatment with Etomoxir does not reduce the degree of noise-induced hearing loss from a 2- hour 112 dB SPL, 8-16 kHz exposure. 24 hours after the exposure, mice were treated with a single dose of Etomoxir (35 mg/kg IP) or saline (vehicle). ABR and DPOAEs were recorded on the last day of treatment (baseline), and on days 1 (after Etomoxir treatment), 7, and 14 after noise. Higher thresholds represent reduced hearing ability. Data are shown as mean ± SEM and were evaluated at each time point by mixed model 2-way ANOVA followed by Šídák multiple- comparison correction. Saline, n=8 mice; Etomoxir, n=8 mice. ns = not significant.

**Supplemental Figure 4:**
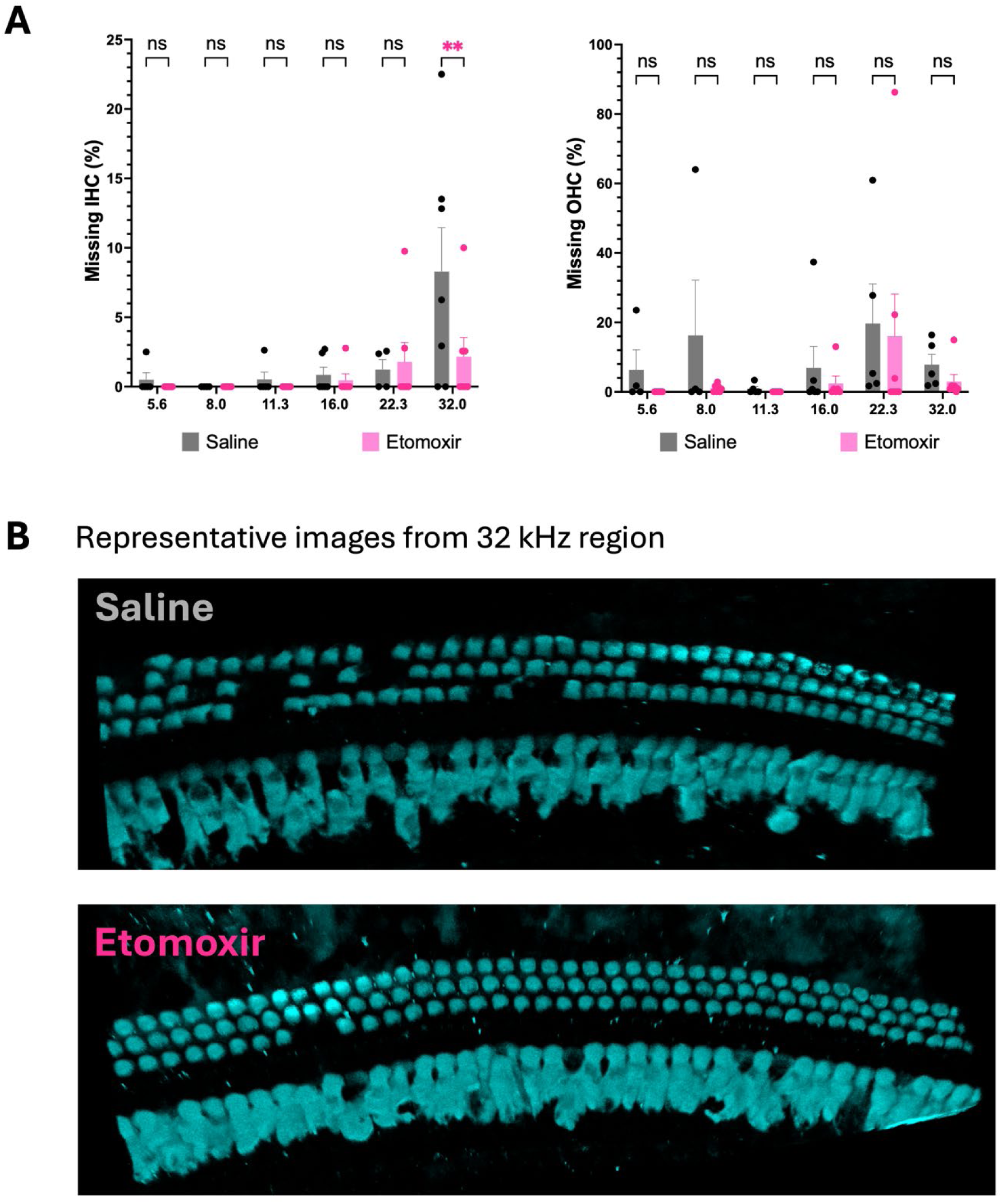
Prophylactic treatment with Etomoxir reduces losses of inner hair cells (IHCs) in the cochlear basal turn. Mice were treated daily with Etomoxir (35 mg/kg IP) or saline (vehicle) for 16 days before 112 dB SPL noise exposure. Samples were harvested ∼1 week after the last ABR or DPOAE recording and stained for MyoVIIa. Data are shown as mean ± SEM and were evaluated at each time point by mixed model 2-way ANOVA followed by Šídák multiple-comparison correction. Etomoxir, n=7 mice; Saline, n=7 mice. ns = not significant; **p < 0.01.

## References

1. Natarajan, N., Batts, S., and Stankovic, K.M. (2023). Noise-Induced Hearing Loss. Journal of Clinical Medicine 12, 2347. 10.3390/jcm12062347.

2. Kohrman, D., Wan, G., Cassinotti, L., and Corfas, G. (2020). Hidden Hearing Loss: A Disorder with Multiple Etiologies and Mechanisms. Cold Spring Harbor Perspectives in Medicine 10, a035493. 10.1101/cshperspect.a035493.

3. Ji, L., Borges, B.C., Martel, D.T., Wu, C., Liberman, M.C., Shore, S.E., and Corfas, G. (2024). From hidden hearing loss to supranormal auditory processing by neurotrophin 3- mediated modulation of inner hair cell synapse density. PLoS Biol 22, e3002665. 10.1371/journal.pbio.3002665.

4. Wang, Y., Hirose, K., Liberman, M.C., Wang, Y., Hirose, K., and Liberman, M.C. (2002). Dynamics of Noise-Induced Cellular Injury and Repair in the Mouse Cochlea. Journal of the Association for Research in Otolaryngology 2002 3:3 *3*. 10.1007/s101620020028.

5. Ji, L., Lee, H.-J., Wan, G., Wang, G.-P., Zhang, L., Sajjakulnukit, P., Schacht, J., Lyssiotis, C.A., and Corfas, G. (2019). Auditory metabolomics, an approach to identify acute molecular effects of noise trauma. Scientific Reports 9. 10.1038/s41598-019-45385-8.

6. Jung, W., Kim, J., Cho, I.Y., Jeon, K.H., and Song, Y.-M. (2022). Association between Serum Lipid Levels and Sensorineural Hearing Loss in Korean Adult Population. Korean Journal of Family Medicine 43, 334–343. 10.4082/kjfm.21.0148.

7. Evans, M.B., Tonini, R., Shope, C.D., Oghalai, J.S., Jerger, J.F., Insull, W., and Brownell, W.E. (2006). Dyslipidemia and Auditory Function. Otology & Neurotology 27, 609–614. 10.1097/01.mao.0000226286.19295.34.

8. Tan, H.E., Lan, N.S.R., Knuiman, M.W., Divitini, M.L., Swanepoel, D.W., Hunter, M., Brennan-Jones, C.G., Hung, J., Eikelboom, R.H., and Santa Maria, P.L. (2018). Associations between cardiovascular disease and its risk factors with hearing loss—A cross-sectional analysis. Clinical Otolaryngology 43, 172–181. 10.1111/coa.12936.

9. Chang, N.C., Yu, M.L., Ho, K.Y., and Ho, C.K. (2007). Hyperlipidemia in noise-induced hearing loss. Otolaryngology–Head and Neck Surgery 137, 603–606. 10.1016/j.otohns.2007.04.022.

10. Doosti, A., Lotfi, Y., and Bakhshi, E. (2016). Effects of Hyperlipidemia on Noise Induced Hearing Loss (NIHL). Indian Journal of Otolaryngology and Head & Neck Surgery 68, 211–213. 10.1007/s12070-015-0855-2.

11. Park, J.S., Kim, S.W., Park, K., Choung, Y.H., Jou, I., and Park, S.M. (2012). Pravastatin attenuates noise-induced cochlear injury in mice. Neuroscience 208, 123–132. 10.1016/j.neuroscience.2012.02.010.

12. Cockcroft, S. (2021). Mammalian lipids: structure, synthesis and function. Essays in Biochemistry 65, 813–845. 10.1042/ebc20200067.

13. Harayama, T., and Riezman, H. (2018). Understanding the diversity of membrane lipid composition. Nature Reviews Molecular Cell Biology 19, 281–296. 10.1038/nrm.2017.138.

14. Antonny, B., Vanni, S., Shindou, H., and Ferreira, T. (2015). From zero to six double bonds: phospholipid unsaturation and organelle function. Trends in Cell Biology 25, 427–436. 10.1016/j.tcb.2015.03.004.

15. Canlon, B., and Schacht, J. (1983). Acoustic stimulation alters deoxyglucose uptake in the mouse cochlea and inferior colliculus. Hearing Research 10, 217–226. 10.1016/0378-5955(83)90055-2.

16. 16. Schacht, J. (1986). Biochemistry of Cochlear Function and pathology. In 01. (Copyright© 1986 by Thieme Medical Publishers, Inc.), pp. 101-114.

17. Houten, S.M., Violante, S., Ventura, F.V., and Wanders, R.J.A. (2016). The Biochemistry and Physiology of Mitochondrial Fatty Acid β-Oxidation and Its Genetic Disorders. Annual Review of Physiology 78, 23–44. 10.1146/annurev-physiol-021115-105045.

18. O’Connor, R.S., Guo, L., Ghassemi, S., Snyder, N.W., Worth, A.J., Weng, L., Kam, Y., Philipson, B., Trefely, S., Nunez-Cruz, S., et al. (2018). The CPT1a inhibitor, etomoxir induces severe oxidative stress at commonly used concentrations. Scientific Reports 8. 10.1038/s41598-018-24676-6.

19. Ceccarelli, S.M., Chomienne, O., Gubler, M., and Arduini, A. (2011). Carnitine palmitoyltransferase (CPT) modulators: a medicinal chemistry perspective on 35 years of research. Journal of medicinal chemistry 54, 3109–3152.

20. Christian, Rohrbach, M., Karrasch, M., Boehm, E., Polonski, L., Ponikowski, P., and Rhein, S. (2007). A double-blind randomized multicentre clinical trial to evaluate the efficacy and safety of two doses of etomoxir in comparison with placebo in patients with moderate congestive heart failure: the ERGO (etomoxir for the recovery of glucose oxidation) stud. Clinical Science 113, 205–212. 10.1042/cs20060307.

21. Puschner, B., and Schacht, J. (1997). Energy metabolism in cochlear outer hair cells in vitro. Hearing Research 114, 102–106. 10.1016/s0378-5955(97)00163-9.

22. Wang, P., Zhang, F., Pan, L., Tan, Y., Song, F., Ge, Q., Huang, Z., and Yao, L. (2021). Inhibiting Cardiac Mitochondrial Fatty Acid Oxidation Attenuates Myocardial Injury in a Rat Model of Cardiac Arrest. Oxidative Medicine and Cellular Longevity 2021, 1–11. 10.1155/2021/6622232.

23. Rosca, M.G., Vazquez, E.J., Chen, Q., Kerner, J., Kern, T.S., and Hoppel, C.L. (2012). Oxidation of Fatty Acids Is the Source of Increased Mitochondrial Reactive Oxygen Species Production in Kidney Cortical Tubules in Early Diabetes. Diabetes 61, 2074–2083. 10.2337/db11-1437.

24. Oishi, N., and Schacht, J. (2011). Emerging treatments for noise-induced hearing loss. Expert Opinion on Emerging Drugs 16, 235–245. 10.1517/14728214.2011.552427.

25. Tan, W.J.T., and Song, L. (2023). Role of mitochondrial dysfunction and oxidative stress in sensorineural hearing loss. Hearing Research 434, 108783. 10.1016/j.heares.2023.108783.

26. Wang, C., Qiu, J., Li, G., Wang, J., Liu, D., Chen, L., Song, X., Cui, L., and Sun, Y. (2022). Application and prospect of quasi-targeted metabolomics in age-related hearing loss. Hearing Research 424, 108604. 10.1016/j.heares.2022.108604.

27. Wan, H., Wang, W., Liu, J., Zhang, Y., Yang, B., Hua, R., Chen, H., Chen, S., and Hua, Q. (2023). Cochlear metabolomics, highlighting novel insights of purine metabolic alterations in age-related hearing loss. Hearing Research 440, 108913.

28. 10.1016/j.heares.2023.108913.

28. Miao, L., Zhang, J., Yin, L., and Pu, Y. (2022). Metabolomics Analysis Reveals Alterations in Cochlear Metabolic Profiling in Mice with Noise-Induced Hearing Loss. BioMed Research International 2022, 1–15. 10.1155/2022/9548316.

29. Dixon, M., Stafford, J., White, F., Clayton, N., and Gallagher, J. (2014). Disorders of mitochondrial energy metabolism, lipid metabolism and other disorders. Clinical Paediatric Dietetics, 588-636.

31. 30. Wolfe, L., Jethva, R., Oglesbee, D., and Vockley, J. (2018). Short-chain acyl-CoA dehydrogenase deficiency.

31. Suzuki, J., Hemmi, T., Maekawa, M., Watanabe, M., Inada, H., Ikushima, H., Oishi, T., Ikeda, R., Honkura, Y., Kagawa, Y., et al. (2023). Fatty acid binding protein type 7 deficiency preserves auditory function in noise-exposed mice. Scientific Reports 13. 10.1038/s41598-023-48702-4.

32. Miwa, T., Tarui, A., Kouga, T., Asai, Y., Ogita, H., Fujikawa, T., and Hakuba, N. (2025). N (1)-methylnicotinamide promotes age-related cochlear damage via the overexpression of SIRT1. Front Cell Neurosci 19, 1542164. 10.3389/fncel.2025.1542164.

33. Fujita, T., Yamashita, D., Irino, Y., Kitamoto, J., Fukuda, Y., Inokuchi, G., Hasegawa, S., Otsuki, N., Yoshida, M., and Nibu, K.-I. (2015). Metabolomic profiling in inner ear fluid by gas chromatography/mass spectrometry in guinea pig cochlea. Neuroscience Letters 606, 188–193. 10.1016/j.neulet.2015.09.001.

34. Pirttilä, K., Videhult Pierre, P., Haglöf, J., Engskog, M., Hedeland, M., Laurell, G., Arvidsson, T., and Pettersson, C. (2019). An LCMS-based untargeted metabolomics protocol for cochlear perilymph: highlighting metabolic effects of hydrogen gas on the inner ear of noise exposed Guinea pigs. Metabolomics 15. 10.1007/s11306-019-1595-1.

35. Qi, G., Jinge, T., Handai, Q., Runnan, H., Qingqing, J., Ning, Y., Shiming, Y., and and Han, D. (2025). Metabolome modification and underlying biomarker of noise-induced hearing loss Guinea pig cochlear fluid. Acta Oto-Laryngologica 145, 101–114. 10.1080/00016489.2024.2445738.

36. Kopke, R.D., Coleman, J.K.M., Liu, J., Campbell, K.C.M., and Riffenburgh, R.H. (2002). Enhancing Intrinsic Cochlear Stress Defenses to Reduce Noise-Induced Hearing Loss. The Laryngoscope 112, 1515–1532. 10.1097/00005537-200209000-00001.

37. Henderson, D., Bielefeld, E.C., Harris, K.C., and Hu, B.H. (2006). The Role of Oxidative Stress in Noise-Induced Hearing Loss. Ear & Hearing 27, 1–19. 10.1097/01.aud.0000191942.36672.f3.

38. Coleman, J.K.M., Kopke, R.D., Liu, J., Ge, X., Harper, E.A., Jones, G.E., Cater, T.L., and Jackson, R.L. (2007). Pharmacological rescue of noise induced hearing loss using N-acetylcysteine and acetyl-l-carnitine. Hearing Research 226, 104–113. 10.1016/j.heares.2006.08.008.

39. Choi, C.H., Chen, K., Du, X., Floyd, R.A., and Kopke, R.D. (2011). Effects of delayed and extended antioxidant treatment on acute acoustic trauma. Free Radic Res 45, 1162–1172. 10.3109/10715762.2011.605360.

40. Rhee, C.-K., and Chang, S.-Y. (2021). Combination photobiomodulation/N-acetyl-L-cysteine treatment appears to mitigate hair cell loss associated with noise-induced hearing loss in rats. Lasers in Medical Science 36, 1941–1947. 10.1007/s10103-021-03304-2.

41. Ohinata, Y., Miller, J.M., Altschuler, R.A., and Schacht, J. (2000). Intense noise induces formation of vasoactive lipid peroxidation products in the cochlea. Brain Research 878, 163–173. 10.1016/s0006-8993(00)02733-5.

42. Tanaka, S., Tabuchi, K., Hoshino, T., Murashita, H., Tsuji, S., and Hara, A. (2010). Protective effects of exogenous GM-1 ganglioside on acoustic injury of the mouse cochlea. Neuroscience Letters 473, 237–241. 10.1016/j.neulet.2010.02.057.

43. Teraoka, M., Hato, N., Inufusa, H., and You, F. (2024). Role of Oxidative Stress in Sensorineural Hearing Loss. International Journal of Molecular Sciences 25, 4146. 10.3390/ijms25084146.

45. 44. Maniaci, A., La Via, L., Lechien, J.R., Sangiorgio, G., Iannella, G., Magliulo, G., Pace, A., Mat, Q., Lavalle, S., and Lentini, M. (2024). Hearing Loss and Oxidative Stress: A Comprehensive Review. Antioxidants 13, 842. 10.3390/antiox13070842.

45. Henderson, D., Bielefeld, E.C., Harris, K.C., and Hu, B.H. (2006). The Role of Oxidative Stress in Noise-Induced Hearing Loss. Ear and hearing 27. 10.1097/01.aud.0000191942.36672.f3.

46. Zhou, Y., Fang, C., Yuan, L., Guo, M., Xu, X., Shao, A., Zhang, A., and Zhou, D. (2023). Redox Homeostasis Dysregulation in Noise-Induced Hearing Loss: Oxidative Stress and Antioxidant Treatment. Journal of Otolaryngology - Head & Neck Surgery 52. 10.1186/s40463-023-00686-x.

48. 47. Le Prell, C.G. (2022). Prevention of Noise-Induced Hearing Loss Using Investigational Medicines for the Inner Ear: Previous Trial Outcomes Should Inform Future Trial Design. Antioxidants & Redox Signaling 36, 1171–1202. 10.1089/ars.2021.0166.

48. Bligh, E.G., and Dyer, W.J. (1959). A rapid method of total lipid extraction and purification. Can J Biochem Physiol 37, 911–917. 10.1139/o59-099.

49. Gika, H.G., Macpherson, E., Theodoridis, G.A., and Wilson, I.D. (2008). Evaluation of the repeatability of ultra-performance liquid chromatography-TOF-MS for global metabolic profiling of human urine samples. J Chromatogr B Analyt Technol Biomed Life Sci 871, 299–305. 10.1016/j.jchromb.2008.05.048.

50. Chambers, M.C., Maclean, B., Burke, R., Amodei, D., Ruderman, D.L., Neumann, S., Gatto, L., Fischer, B., Pratt, B., Egertson, J., et al. (2012). A cross-platform toolkit for mass spectrometry and proteomics. Nat Biotechnol 30, 918–920. 10.1038/nbt.2377.

51. Kind, T., Meissen, J.K., Yang, D., Nocito, F., Vaniya, A., Cheng, Y.S., Vandergheynst, J.S., and Fiehn, O. (2012). Qualitative analysis of algal secretions with multiple mass spectrometric platforms. J Chromatogr A 1244, 139–147. 10.1016/j.chroma.2012.04.074.

52. Meissen, J.K., Yuen, B.T., Kind, T., Riggs, J.W., Barupal, D.K., Knoepfler, P.S., and Fiehn, O. (2012). Induced pluripotent stem cells show metabolomic differences to embryonic stem cells in polyunsaturated phosphatidylcholines and primary metabolism. PLoS One 7, e46770. 10.1371/journal.pone.0046770.

53. Ejsing, C.S., Duchoslav, E., Sampaio, J., Simons, K., Bonner, R., Thiele, C., Ekroos, K., and Shevchenko, A. (2006). Automated identification and quantification of glycerophospholipid molecular species by multiple precursor ion scanning. Anal Chem 78, 6202–6214. 10.1021/ac060545x.

54. Ji, L., Borges, B.C., Martel, D.T., Wu, C., Liberman, M.C., Shore, S.E., and Corfas, G. (2024). From hidden hearing loss to supranormal auditory processing by neurotrophin 3- mediated modulation of inner hair cell synapse density. PLOS Biology 22, e3002665. 10.1371/journal.pbio.3002665.

